# Structure-aware Protein Solubility Prediction From Sequence Through Graph Convolutional Network And Predicted Contact Map

**DOI:** 10.1101/2020.06.24.169011

**Authors:** Jianwen Chen, Shuangjia Zheng, Huiying Zhao, Yuedong Yang

## Abstract

**Motivation:** Protein solubility is significant in producing new soluble proteins that can reduce the cost of biocatalysts or therapeutic agents. Therefore, a computational model is highly desired to accurately predict protein solubility from the amino acid sequence. Many methods have been developed, but they are mostly based on the one-dimensional embedding of amino acids that is limited to catch spatially structural information.

**Results:** In this study, we have developed a new structure-aware method to predict protein solubility by attentive graph convolutional network (GCN), where the protein topology attribute graph was constructed through predicted contact maps from the sequence. GraphSol was shown to substantially out-perform other sequence-based methods. The model was proven to be stable by consistent R^2^ of 0.48 in both the cross-validation and independent test of the *eSOL* dataset. To our best knowledge, this is the first study to utilize the GCN for sequence-based predictions. More importantly, this architecture could be extended to other protein prediction tasks.

**Availability:** The package is available at http://biomed.nscc-gz.cn

**Supplementary information:** Supplementary data are available at *Bioinformatics* online.

## 1 Introduction

Over the past 20 years, recombinant protein had played a vital role in biotechnology and medicine, including novel therapeutic protein drugs and antibodies (Habibi, et al., 2014). Recombinant proteins are mostly produced by genetic engineering in *Escherichia coli (E*.*coli)* (Chan, et al., 2010). However, low solubility and activity of proteins expressed by *E*.*coli* limited the production efficiency even though the standard workflow and logical strategies have been widely deployed in biopharmaceutical industries. According to statistics, over 30% of recombinant proteins are not soluble (Samak, et al., 2012), 33∼35% of all expressed non-membrane proteins are insoluble, and 25∼57% of soluble proteins are prone to aggregate at higher concentrations (Fang and Fang, 2013). Moreover, the heterologous expression often suffers from low levels of production and insoluble recombinant proteins forming inclusion bodies. Therefore, the solubility of proteins plays an important role in the production of proteins for the biotechnological and pharmaceutical industries.

To enhance the performance of recombinant proteins, many experimental technologies have been developed, e.g. directed evolution, immobilization, designing better promoters, optimizing codon usage, and changing culture conditions including media and temperature (Agostini, et al., 2012; Madhavan, et al., 2017). However, such empirical optimizations are labor-intensive and time-consuming. A precise computational model is highly desired so that protein solubility can be effectively predicted. Theoretically, given an exact experimental condition (i.e. temperature, expression host, etc.), the solubility is determined only by its primary structure that is decided by the sequence (Smialowski, et al., 2012). To this end, two types of computational approaches have been proposed to predict the protein solubility: physical-based and machine/deep learning-based methods.

In terms of the physical-based techniques, most works focused on making use of extensive molecular dynamics simulations to evaluate the free energy difference between aggregation and solution phases. However, these methods are usually of limited accuracy due to difficulties in evaluating the conformational entropy and solvent contributions. Furthermore, these atom-level methods are sensitive to structural fluctuations and can’t process protein flexibility well (Hou, et al., 2020).

For the machine/deep learning techniques, several sequence-based methods have been developed for protein solubility prediction including *PROSO II* (Smialowski, et al., 2012), *CCSOL* (Agostini, et al., 2012), *SOLpro* (Magnan, et al., 2009), and the scoring card method (*SCM*) (Huang, et al., 2012). The majority of these methods adopted the support vector machine(*SVM*) (AK, 2002) as the core discriminative model on biologically relevant handcrafted features from protein sequences to discriminate the soluble and insoluble proteins. The newly proposed method, *PaRSnIP* (Rawi, et al., 2018) was developed by identifying correlations of protein solubilities positively with fractions of exposed residues while negatively with tri-peptide stretches containing multiple histidines. *Solx-Plain* employed an *XGBoost-based* model (Chen and Guestrin, 2016) by using various features derived from protein sequences (Mall, 2019).

With the development of deep learning techniques, many end-to-end methods have been developed. *DeepSol* built a convolutional neural network to construct non-linear high-dimensional k-mer vector spaces with essential information for predicting protein solubility (Khurana, et al., 2018). *ProGAN* generated extra data from a Generative Adversarial Networks (GAN) (Goodfellow, et al., 2014) that had been learned by the training set to improve the final performance (Han, et al., 2019). However, these methods are mostly based on LSTM and CNN and didn’t utilize spatial information of protein molecules. Though our recent studies indicated that protein structure could be well represented by the residue-pairwise distance matrix through CNN (Chen, et al., 2020; Zheng, et al., 2020), the contacted structural information is only implicitly included that can’t fully utilize the relations between contacted residues.

In the past few years, graph neural network (GNN) was raised to represent the protein structure in various of deep learning-based methods and has made successes (Gligorijević, et al., 2018; Zamora-Resendiz and Crivelli, 2019). However, these methods demand experimentally obtained 3D structures that are hard to acquire for aggregation proteins and thus are not appropriate for sequence-based protein solubility prediction.

With recent developments in protein structure prediction, the prediction of protein contact map has been greatly improved according to the critical assessment of protein structure prediction (CASP) (Schaarschmidt, et al., 2018). For example, *Hanson et al* had developed a novel sequence-based method in predicting protein contact map, which aimed to capture these deep, underlying relationships between residue-residue pairs in spatial dimensions for protein ‘image’ at each layer (Hanson, et al., 2018). Compared to other algorithms, the predicted protein contact map integrates all their advantages so that it can represent 2D structural features directly in high accuracy, enabling the construction of accurate protein graphical representations from protein sequence.

In this study, we purposed a novel structure-aware method *GraphSol* for protein solubility prediction from the sequence by combining predicted contact maps and graph neural networks. The predicted contact maps were employed to construct protein graphs, and the attentive-based graph convolutional network made the predictions through mapping the nodes (amino acids) embedding to the graph full content embedding. We performed our model in the *eSOL* database (Niwa, et al., 2009) and obtained state-of-the-art performance. To the best of our knowledge, this is the first study to make sequence-based predictions for proteins through a graph neural network. Such architecture could be easily applied to extensive tasks on proteins, e.g. protein function prediction, protein folding, and drug design.

## 2 Methods

### 2.1 Overview

In this work, we convert the protein solubility prediction task as a graph-based regression problem. Given a protein sequence that consists of *L* amino acids, the whole protein could thus be expressed as a topological attributed graph *G*(*F, E*), with *F* for the feature set of all residues (nodes) and *E* for the residual contacts (edges) according to predicted protein contact map. Our task aims to learn a mapping function *f*(·) that inputs with predicted residual features and contact map and outputs predicted solubility with continuous scores between [0,1] ∈ ℝ i.e. *f*: *G*(*F, E*) → [0,1]. In this work, *f*(·) is a graph convolutional neural network model that aggerates nodes and edges information on the irregular graph.

### 2.2 Dataset

#### eSOL dataset

To train our model, we employed the *eSOL* dataset from the previous study (Han, et al., 2019). For completeness, we briefly describe the procedure to produce the dataset. The whole solubility database of ensemble *E*.*coli* proteins was downloaded from the *eSOL* website (Niwa, et al., 2009), where the solubility was defined as the ratio of the supernatant fraction to the total fraction in the physiochemical experiments. The 4,132 proteins were mapped to the *NCBI* database by gene names, leading to a total of 3,147 protein sequences. From these sequences, 75% (2363 proteins) were randomly selected as the training set, and the remaining 784 proteins were used as the independent test.

#### S. cerevisiae dataset

For an external independent test, we selected another protein dataset collected by (Hou, et al., 2020) from the *S. cerevisiae*. This dataset was derived by including 108 proteins having corresponding 3D structures. The solubilities were measured also by the cell-free expression called PURE (Shimizu, et al., 2005).

### 2.3 Protein representation

#### 2.3.1 Node features

We devised four groups of protein features that were used to train the GraphSol predictor model.

##### Blosum62

Instead of simple one-hot encoding, we have encoded residues by Blosum62 (Mount, 2008), which is a widely used 20 × 20 matrix for substitutions between 20 standard amino acid types according to alignments of homologous protein sequences.

##### Physicochemical Properties

We utilized a set of 7 physicochemical properties for amino acid types (AAPHY7) (Meiler, et al., 2001). These features include steric parameters, hydrophobicity, volume, polarizability, isoelectric point, helix probability, and sheet probability.

##### Evolutionary information

Evolutionarily conserved residues may contain the motifs related to protein properties (such as solubility) in biological sequences (Narjeskhatoon Habibi1* and 2014). Here, we employed the position-specific scoring matrix (PSSM) and the HHM matrix. To be specific, the PSSM profile was produced by PSI-BLAST v2.7.1 (Altschul, et al., 1997) with the UniRef90 sequence database after 3 iterations. The HHM profile was produced by HHblits v3.0.3 in aligning the UniClust30 profile HHM database (Mirdita, et al., 2017) with default parameters.

##### Predicted Structural properties

Predicted structural properties are highly related to solubility in the previous study (Khurana, et al., 2018). Herein, we derived the predicted structural features from SPIDER3 (Heffernan, et al., 2017), one of the most accurate predictors. The feature group includes 14 features: 1) three probability values respectively for three secondary structure states, 2) Relative Solvent-Accessible Surface Area (ASA), 3) eight values for the sine/cosine values of backbone torsion angles (phi, psi, theta, tau), and 4) Half-Sphere Exposures based on the C_α_ atom (HSE-up and HSE-down).

Finally, these feature groups constructed the node feature matrix X ∈ ℝ^*L*×94^ with *L* representing the length of a protein sequence. **Table 1** listed all node feature groups with their dimensions. All data were standardized to zero mean and unit variance before input into the neural network.

**Table 1.** Node features and dimensions

| Group | Node Features | Names | Dimensions |
| --- | --- | --- | --- |
| 1 | BLOSUM matrix | BLOSUM62 | 20 |
| 2 | Physicochemical Properties | AAPHY7 | 7 |
| 3 | Position-Specific Scoring Matrix | PSSM | 20 |
|  | Hidden Markov Matrix | HHM | 30 |
| 4 | Structural Properties predicted by SPIDER3 | SPIDER3 | 14 |

#### 2.3.2 Edge features

In order to construct the edges for the protein attribute graph representation, we make predictions of the protein contact map from a sequence by *SPOT-Contact* (Hanson, et al., 2018), which outputs the possibilities to form contacts between all residue pairs in one protein. In default, the graph is a fully connected graph constructed with each edge valued as the predicted contact probability of the corresponding residue pair. As the actual number of contacts in a protein is approximately proportional to the length, we also test constructing the protein attribute graph by setting up edges for α × L residual pairs with the highest predicted contact probability, as also used in the critical assessment of protein structure prediction (CASP) (Schaarschmidt, et al., 2018). The α = 1∼7 was used as suggested by (Emerson and Amala, 2017). Herein, we tested two schemes to construct each selected edge by setting the value as “1” (discrete) and the predicted contact probability (continuous), respectively. The neighbored residues in a sequence are always connected. Notably, though the fully connected graph (default mode) was shown to perform the best according to our results, the partial edges decrease the computational and memory complexity from O(L^2^) to O(L).

### 2.4 Deep Learning Framework

Our graph-based model consists of three parts. As shown in **Figure 1**, the first part is a graph convolution network (GCN), which aggregates protein structural information from its nodes and edges during iterations. The second part is a self-attention pooling layer, which transforms the node hidden state of varied sizes to the graph representation vector with a fixed size. Finally, this fix-sized vector goes through full connection layers to predict the protein solubility.

**Figure 1.**
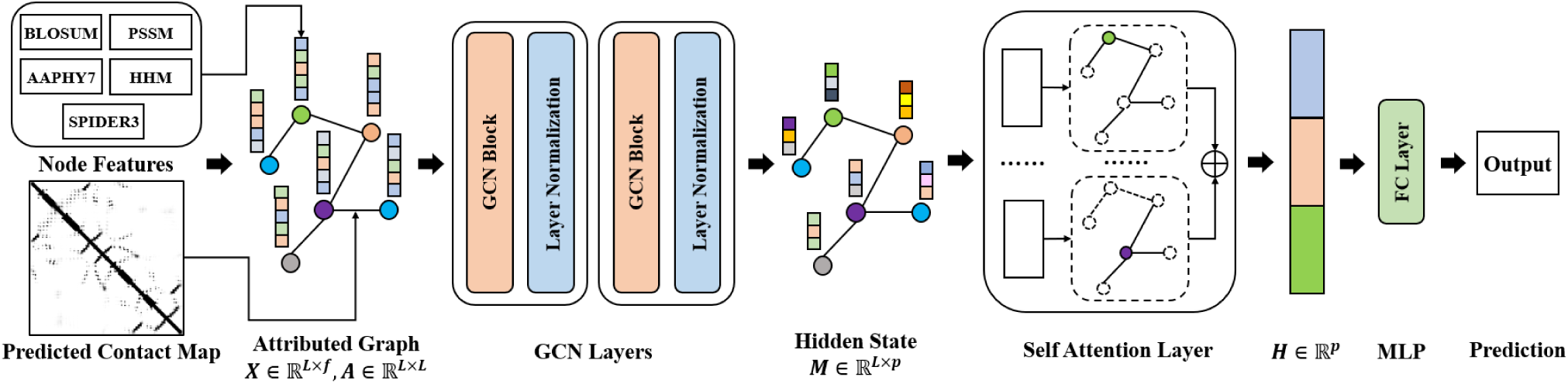
The overall framework of the GraphSol model. The compiled node features **(X)** and contact maps **(A)** were utilized to construct the GCN networks, which were convoluted by two GCN blocks and layer normalizations for hidden states **(M)**. The hidden states were converted by the self-attentive layer as a fixed length of the vector **(H)**, which were input to the MLP to make the final prediction.

#### 2.4.1 Graph Convolution Network

Given a protein sequence with *L* amino acids, the protein is represented by the feature matrix *X* ∈ ℝ^*L*×*f*^ for nodes and contact matrix *A* ∈ ℝ^*L*×*L*^ for edges with *f* as the number of features for nodes. Our graph convolution network (GCN) takes the following formula (Kipf and Welling, 2016):

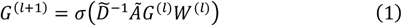

where *Ã* = *A* + *I*_*L*_ is the adjacency matrix by adding the edge matrix *A* determined by predicted contact map and the identity matrix *I*_*L*_ for self-loops. 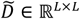 is a diagonal degree matrix with 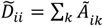 that is used to normalize *Ã* to sum up to 1.0 in each row. *G*^(*l*)^ ∈ ℝ^*L*×*f*^ is the activation hidden matrix in the *l*^*th*^ layers with the initial state 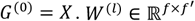 is a weight matrix of layer-specific trainable parameters to map the iteration to a lower dimension rich-information space with a size of *f*′. σ denotes a nonlinear activation function and we use the ReLU(·) function here. After each GCN layer, a normalization layer is added to rescale the output to [0,1], which was found to accelerate the converge of the GCN layers. The final output of the last GCN layer is integrated as

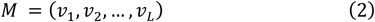

where *v*_*i*_ is a *p* dimensional vector token embedding for the *i*^*th*^ node. As a result, *M* is a 2D matrix to integrate all token embeddings with ℝ^*L*×*p*^.

#### 2.4.2 Self-Attention pooling

Note that the output matrix M is dependent on the protein length, which is a variable scale. To obtain a fixed size of protein representation, a readout transformation is essential to eliminate the size variance and sequence permutation variance (Zheng, et al., 2019). Herein, we employ the self-attention mechanism (Lin, et al., 2017), which computes the weight coefficients *T* ∈ ℝ^*r*×*L*^ with *r* for the number of attention groups by:

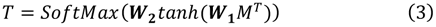

where *M*^*T*^ is the transposition of *M* ∈ ℝ^*L*×*p*^. ***W***_**1**_ ∈ ℝ^*q*×*p*^ and ***W***_**2**_ ∈ ℝ^*r*×*q*^ are two learned attention matrices with the hyper-parameter *q*. The *SoftMax* function standardizes each row of the computed weights, to sum up to 1. Intuitively, the *r* groups of attention coefficients of *A* assess the associations of each residue with the solubility from different views. Thus, we extract the overall features by multiplying T and M, and average all *r* groups of attention coefficients for the graph representation *H* ∈ ℝ^1×*p*^ by

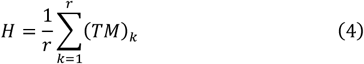

#### 2.4.3 Multilayer Perceptron

The output of self-attention pooling was input to the Multilayer Perceptron (MLP) to predict the solubility *S* by

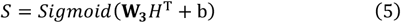

where **W**_**3**_ ∈ ℝ^1×p^ is the weight matrix and b ∈ ℝ is the bias item. The sigmoid function maps the value to (0,1) for solubility prediction.

### 2.5 Training and Evaluation

#### 2.5.1 Hyper-parameter tuning

Our model for solubility prediction includes multiple hyperparameters. We tested crucial hyperparameters and the range of values as follows:

1. **GCN Layers:** A higher number of GCN layers means the wider and deeper information aggregated from the edge and node features. On the other hand, excessive layers would cause a decrease in final predicted accuracy due to vanishing gradients. Therefore, it is crucial to keep a balance between the layers and the algorithm complexity. We test the following settings {1,2,3,4} and found 2 layers to be the optimal value after tuning on the validation sets.
2. **GCN Middle Dimensions:** These hyper-parameters control the channel dimensions in all stacking GCN layers including the final GCN layer. These hyper-parameters affect the matrices that are transferred into self-attention pooling to identify the key soluble fragment of the protein. Therefore, we should construct a suitable size for matrices for liberating the rich-information regions as well as the distinguishability of different proteins. As a result, the optimal parameters are 64 dimensions for the last layer, and 256 for others.
3. **Attention Heads:** The attention heads provide weight coefficients to focus on key residues for the solubility prediction, and different heads enable the attention of multiple regions from different views. We tested the number of attention heads from 1 to 10 and found 4 attention heads provided the best performance in the validations.

In addition, the models were trained for different epochs using Adam optimizer (Kingma and Ba, 2014). **Table S1** showed the optimal hyper-parameters by the grid search.

#### 2.5.2 Cross-Validation and Independent Test

We performed the 5-fold cross-validation on the training set of the *eSOL* dataset. That is, proteins in the training set were separated into five parts (folds). In each round four folds were employed to train a model that was evaluated on the left one-fold. This process was repeated for 5 times, and the performances of five predictions were averaged as the validation performance. To reduce fluctuations by the random splitting of five-folds, we have split the training set with five random seeds and took an average of final performances. The validations were used to optimize all hyper-parameters. With the optimal hyper-parameters, a model was trained by all training set and independently tested on the two independent test sets.

#### 2.5.3 Evaluation indicators

The neural network was trained to minimize the root of mean squared error (RMSE), and the coefficient of determination (R^2^) was used to evaluate our models and optimize the hyper-parameters. Since many compared methods (Agostini, et al., 2012; Hebditch, et al., 2017; Khurana, et al., 2018; Rawi, et al., 2018) have been developed to classify whether a protein is soluble or not, we also separated all proteins into two classes (soluble or not) by a threshold of 0.5 for the predicted and actual solubility. By the definition, the models were evaluated by the area under the Receiver Operating Characteristic *(ROC)* curve (AUC), accuracy, precision, recall, and F1-score defined as:

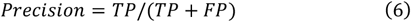

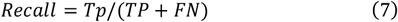

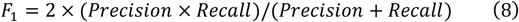

where TP, FP, FN, and FN denote the numbers of true positives (soluble proteins), false positives (non-soluble protein predicted as soluble), true negatives, and false negatives, respectively.

## 3 Results and discussion

### 3.1 Performances on the 5-fold cross-validation and independent test

We investigated the performance of the GraphSol model on the *eSOL* dataset. As shown in **Table 2**, We obtained R^2^ values of 0.476±0.014 and 0.483 for the 5-fold CV and independent test, respectively. When separating the dataset into two discrete states (soluble or not) by a cutoff of 0.5, the AUC values are 0.855 and 0.866 for the 5-fold CV and independent tests, respectively **(Fig S1)**. The similar results by the CV and independent test indicated the robustness of our GraphSol model.

**Table 2.** The R^2^ between the actual solubility scores and those predicted by GraphSol based on individual feature groups, removing each feature group from the final GraphSol model, and recursively adding feature groups according to their importance, respectively.

| Feature groups<sup>a</sup> | CV<sup>d</sup> | Ind. test | Feature groups<sup>b</sup> | CV<sup>d</sup> | Ind. test | Features groups<sup>c</sup> | CV<sup>d</sup> | Ind. test |
| --- | --- | --- | --- | --- | --- | --- | --- | --- |
| - | - | - | GraphSol | **0.476±0.014** | **0.483** | - | - | - |
| BLOSUM | 0.329±0.016 | 0.317 | -BLOSUM | 0.460±0.011 | 0.465 | BLOSUM | 0.329±0.016 | 0.317 |
| AAPHY7 | 0.293±0.014 | 0.289 | -AAPHY7 | 0.465±0.012 | 0.479 | +SPIDER3 | 0.413±0.012 | 0.409 |
| PSSM | 0.333±0.012 | 0.332 | -PSSM | 0.457±0.017 | 0.467 | +PSSM | 0.456±0.011 | 0.453 |
| HHM | 0.337±0.015 | 0.341 | -HHM | 0.455±0.016 | 0.458 | +HHM | 0.465±0.012 | 0.479 |
| SPIDER3 | 0.231±0.019 | 0.227 | -SPIDER3 | 0.428±0.018 | 0.449 | +AAPHY7 | **0.476±0.014** | **0.483** |

<sup>a</sup>Performances based on individual feature groups, <sup>b</sup> by removing each feature group from all feature groups, and <sup>c</sup> by adding feature groups recursively.
<sup>d</sup>Performances by the 5-fold cross-validation.

In order to indicate the importance of feature groups, we assessed the performances by 3 ways in the ablation study. As shown in **Table 2**, when the individual feature group was used as the node features, HHM yielded the highest R^2^ with a value of 0.341 in the independent test. The other evolution-based feature (PSSM) performed similarly but slightly worse than HHM. Not surprisingly, BLOSUM didn’t perform well with R^2^ of 0.317, but better than AAPHY7 features. The predicted structural feature group (SPIDER3) made the worst performance with R^2^ of 0.243. When removing an individual group, on the contrary, the removal of SPIDER3 led to the greatest drop from 0.483 to 0.449. This is likely because SPIDER3 uniquely provided structural information, while other features have supplementary alternatives. Though PSSM and HHM similarly represent evolution information, their removals still caused decreases in performances and generally, HHM is shown more important. The removal of AAPHY7 caused the smallest drop, which is understandable because this feature is a seven-dimensional matrix that is smaller than other feature groups. When we evaluated the model by adding the feature groups recursively, the model showed incremental performances with the addition of each feature group. An interesting fact was that the performance sharply increased from 0.317 to 0.409 after adding SPIDER3 features, which agreed with the relationship between solubility and structural features such as the secondary structure and solvent accessible area.

### 3.2 Evaluating the Impact of Predicted Protein Contact Map

The previous results were based on a fully connected graph with edges weighted according to predicted contact probabilities. As there are a limited number of actual contacts between residue pairs, we tested assigning edges between top α × L residue pairs with the highest predicted contacting probabilities. As shown in **Figure 2**, when not using the predicted contact map (α = 0), i.e. no edges were assigned except between 2-hops neighbored residues, the model achieved R^2^ of 0.462. With the increase of α, the R^2^ has a steady increase followed by a sharp increase from 2 × L to 3 × L with the R^2^ of 0.474. Then a slight but continuous growth was observed with an increase of α. The highest R^2^ of 0.483 was obtained when all residues were connected with the respective predicted probability.

**Figure 2.**
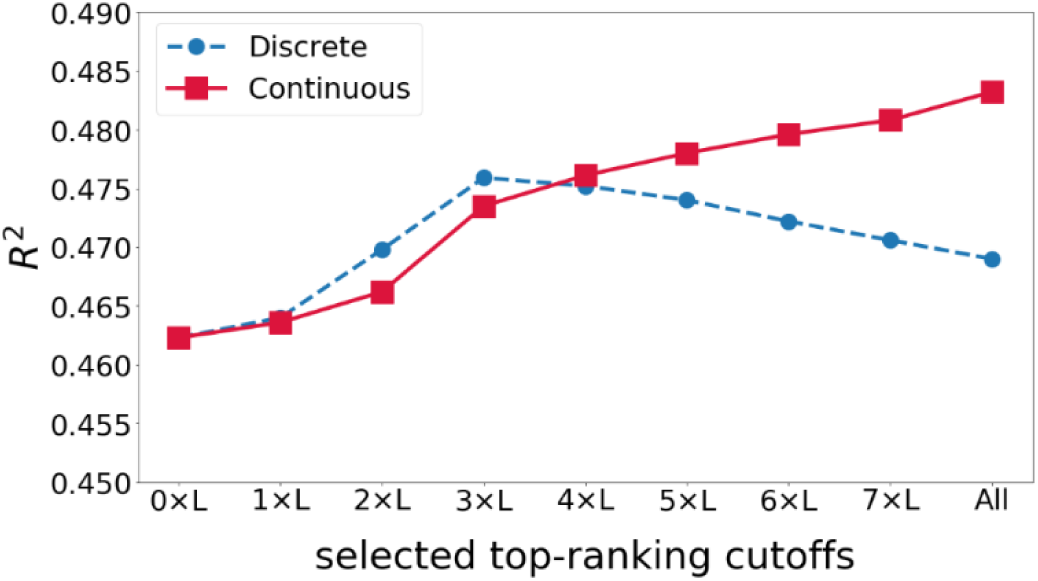
The R^2^ of the GraphSol model changed by selecting the different number of contacts (edges) according to predicted contact maps.

By comparison, we tested the connectivity by discretely assigning all connected edges as 1 with other pairs not connected and labeled to 0. As expected, the R^2^ increased with α from 0 to 3, indicating that the pairs are helpful for the prediction. Afterward, the R^2^ started to decrease likely due to an increase of inaccurately predicted contact pairs. Interestingly, we found there are close R^2^ for the discrete group and continuous group at α nearly to 4.

### 3.3 Comparisons with other methods

Our GraphSol model was compared with state-of-the-art methods. To avoid the impacts caused by different datasets, all machine learning or deep learning-based models were retrained and tested on our training and test set, respectively. As shown in **Table 3**, GraphSol consistently obtained the best results by all measurements as a single method. Even if we didn’t use predicted contact maps, the GraphSol (no-contact) ranked the 2^nd^ that yielded slightly higher *R*^2^ than our self-implemented LSTM model (0.462 vs 0.458). For other methods, *ProGAN* (Han, et al., 2019), a GAN network-based method, achieved the highest *R*^2^ = 0.442, which is 7% lower than our GraphSol (*R*^2^ = 0.483). *DeepSol*, a CNN-based network, achieved *R*^2^of 0.434, which is close to the LSTM. Machine learning techniques didn’t perform well with R^2^ ranging from 0.214 to 0.411. **Figure 3** shows the actual solubility as a function of predicted values by four methods. We found that the deep learning-based methods fitted more accuracy in the region [0.2,0.4], especially the ProGAN and GraphSol model, and GraphSol model performed better in the region of nearly 0.2.

**Table 3.**
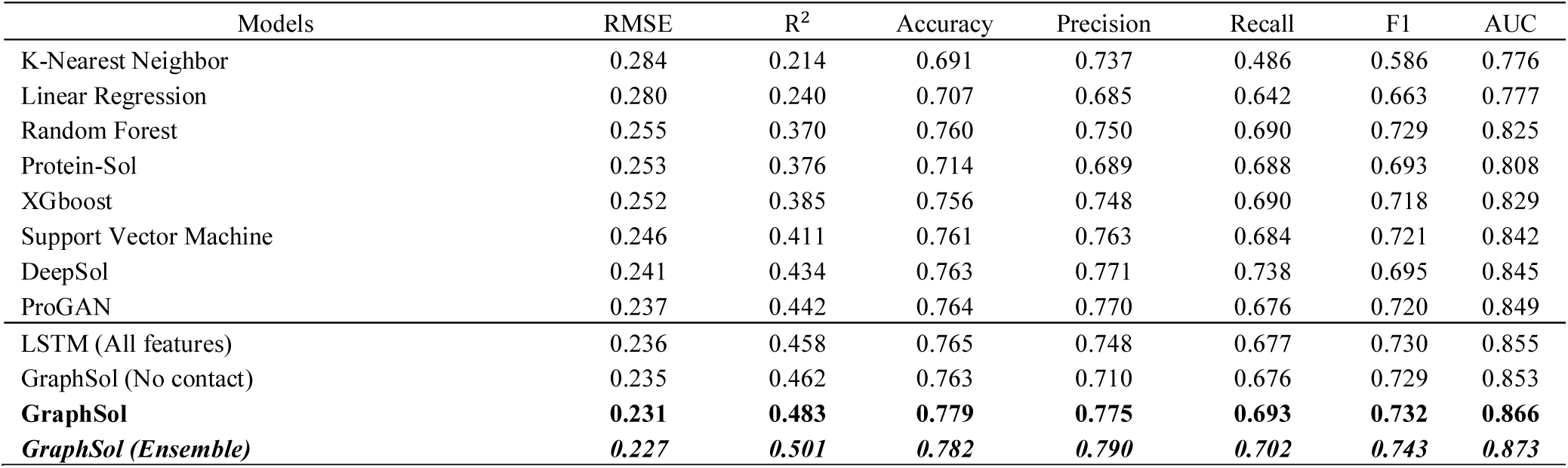
Comparisons of different methods on the eSOL test set.

| Models | RMSE | R<sup>2</sup> | Accuracy | Precision | Recall | F1 | AUC |
| --- | --- | --- | --- | --- | --- | --- | --- |
| K-Nearest Neighbor | 0.284 | 0.214 | 0.691 | 0.737 | 0.486 | 0.586 | 0.776 |
| Linear Regression | 0.280 | 0.240 | 0.707 | 0.685 | 0.642 | 0.663 | 0.777 |
| Random Forest | 0.255 | 0.370 | 0.760 | 0.750 | 0.690 | 0.729 | 0.825 |
| Protein-Sol | 0.253 | 0.376 | 0.714 | 0.689 | 0.688 | 0.693 | 0.808 |
| XGboost | 0.252 | 0.385 | 0.756 | 0.748 | 0.690 | 0.718 | 0.829 |
| Support Vector Machine | 0.246 | 0.411 | 0.761 | 0.763 | 0.684 | 0.721 | 0.842 |
| DeepSol | 0.241 | 0.434 | 0.763 | 0.771 | 0.738 | 0.695 | 0.845 |
| ProGAN | 0.237 | 0.442 | 0.764 | 0.770 | 0.676 | 0.720 | 0.849 |
| LSTM (All features) | 0.236 | 0.458 | 0.765 | 0.748 | 0.677 | 0.730 | 0.855 |
| GraphSol (No contact) | 0.235 | 0.462 | 0.763 | 0.710 | 0.676 | 0.729 | 0.853 |
| **GraphSol** | **0.231** | **0.483** | **0.779** | **0.775** | **0.693** | **0.732** | **0.866** |
| **GraphSol (Ensemble)** | **0.227** | **0.501** | **0.782** | **0.790** | **0.702** | **0.743** | **0.873** |

**Figure 3.**
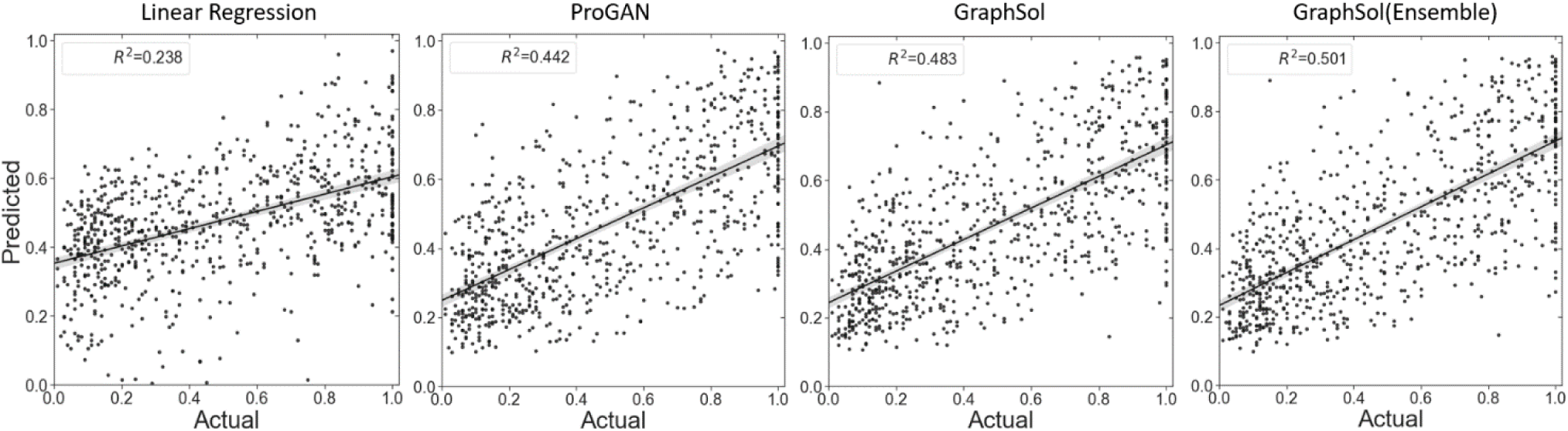
Comparison between actual solubility scores and those predicted by four sequence-based methods on the independent test. The line is the linear least-square fitting to the actual values with the shadow as the 95% confidence intervals for the regressions.

As most of the other methods were designed for predicting discrete states, we also turn the problem into the 2-state classification task. When using a threshold of 0.5 to define soluble proteins or not, our GraphSol achieved the best performances with AUC of 0.866, F-measure of 0.732, and accuracy of 0.779, which are at least 2% better than the best of other methods *ProGAN*. **Figure 4A** compared the ROC curves for six methods, and we can find that the curve of *GraphSol* mostly locates on the top. We also tested how accuracy varies as a function of the threshold which defined a soluble class and the trends were similar **(Figure S2, S3)**.

**Figure 4.**
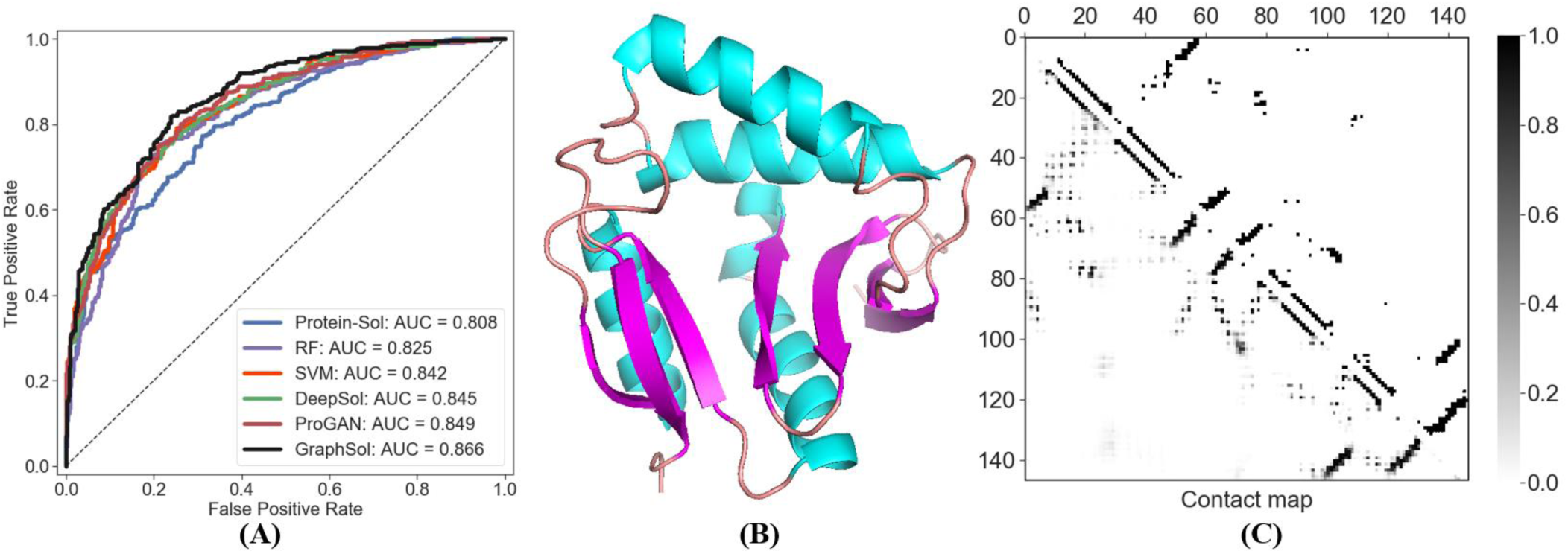
**(A)** Comparisons of the areas under the receiver operating curve (AUC) on the *eSOL* test set, and **(B)** the 3D structure and **(C)** the predicted (left triangular matrix) and actual (the right one) contact map of the selected case (the Peptidyl-lysine N-acetyltransferase *yjaB* gene, PDB ID: 2kcw).

We noticed that GraphSol showed fluctuations in the 5-fold cross-validations, and thus we built an ensemble model by averaging the outputs of 5 trained models during CV on the test set. The *GraphSol (Ensemble)* model was found to further improve the performance by a margin of 3% to 0.501 in R^2^. Other indicators also got varied increases **(Table 3)**.

Moreover, we made comparisons of all methods on the other *S. cerevisiae* test set. Here, we employed the *eSOL* training set to train the model, and have excluded sequences with identity >25% to the *S. cerevisiae* test set. As shown in **Table 4**, GraphSol model yielded R^2^ of 0.358 which is much higher than other sequence-based methods. In comparison, the *DeepSol* and *proGAN* achieved an R^2^ below 0.1. And our ensemble method could further improve *GraphSol* by 3.9%, consistent with the previous results on the *eSOL* dataset. It is noted that all methods achieved lower R^2^ on this dataset. This is likely because this dataset is more challenging, as the *Solart* method was reported to yield R^2^ of only 0.422 even with the use of experimental structures. The *Solart* wasn’t directly compared because most proteins don’t contain experimental structures as also indicated in the *eSOL* dataset.

**Table 4.** Comparisons of different methods on the *S. cerevisiae* test set.

| Predictors | R<sup>2</sup> |
| --- | --- |
| **GraphSol (Ensemble)** | **0.372** |
| **GraphSol** | **0.358** |
| ccSol<sup>a</sup> | 0.302 |
| Protein-Sol<sup>a</sup> | 0.281 |
| CamSol<sup>a</sup> | 0.160 |
| DeepSol<sup>a</sup> | 0.090 |
| ProGAN<sup>b</sup> | 0.084 |

<sup>a</sup>Results produced by (Hou, et al., 2020); <sup>b</sup>Results produced by us.

### 3.4 Case study

To further illustrate our method, we took the Peptidyl-lysine N-acetyl-transferase *yjaB* gene (PDB id: 2kcw) that consisted of 147 residues as an example **(Fig 4B)**. We calculated the C_α_ distance between all residue pairs as the actual contact map. As shown in **Fig 4C**, there are 360 residue pairs with C_α_ distance less than 7.5Å on the actual contact map of the protein. The prediction corresponded to a precision of 0.745 to cover 75.3% actual contacts **(Table S3)**. This high-quality prediction of the residue pairs enabled the accurate construction of the protein attitude graph and the solubility prediction. Besides, when compared to the actual solubility of 0.87, our *GraphSol* model made an accurate prediction of 0.860 and 0.857 by using the continuous and 4 × L discrete predicted contact map, respectively. This similar tendency between **Fig 2** and **Fig S4** also indicated the effectiveness of our method.

## 4 Conclusions

In this study, we introduce the GraphSol model, a sequence-based solubility predictor through graph neural networks. Compared to other methods, we utilized predicted protein contact maps that played a key role in bridging protein topology attribute and attentive graph neural network. We found that the predicted contact probabilities between residues are better to represent the pairwise relations than discrete states. In the future, such a method is potentially useful to protein attribute predictions including protein function prediction and drug design.

## Acknowledgments

This study has been supported by the National Natural Science Foundation of China (61772566, 62041209, and U1611261), Guangdong Key Field R&D Plan (2019B020228001 and 2018B010109006) and Introducing Innovative and Entrepreneurial Teams (2016ZT06D211).

